# Key Issues Review: Evolution on rugged adaptive landscapes

**DOI:** 10.1101/112177

**Authors:** Uri Obolski, Yoav Ram, Lilach Hadany

**Affiliations:** Department of Molecular Biology and Ecology of Plants, Tel-Aviv University, Israel

## Abstract

Adaptive landscapes represent a mapping between genotype and fitness. Rugged adaptive landscapes contain two or more *adaptive peaks:* allele combinations that differ in two or more genes and confer higher fitness than intermediate combinations. How would a population evolve on such rugged landscapes? Evolutionary biologists have struggled with this question since it was first introduced in the 1930’s by Sewall Wright.

Discoveries in the fields of genetics and biochemistry inspired various mathematical models of adaptive landscapes. The development of landscape models led to numerous theoretical studies analyzing evolution on rugged landscapes under different biological conditions. The large body of theoretical work suggests that adaptive landscapes are major determinants of the progress and outcome of evolutionary processes.

Recent technological advances in molecular biology and microbiology allow experimenters to measure adaptive values of large sets of allele combinations and construct *empirical adaptive landscapes* for the first time. Such empirical landscapes have already been generated in bacteria, yeast, viruses, and fungi, and are contributing to new insights about evolution on adaptive landscapes.

In this Key Issues Review we will: (i) introduce the concept of adaptive landscapes; (ii) review the major theoretical studies of evolution on rugged landscapes; (iii) review some of the recently obtained empirical adaptive landscapes; (iv) discuss recent mathematical and statistical analyses motivated by empirical adaptive landscapes, as well as provide the reader with source code and instructions to implement simulations of adaptive landscapes; and (v) discuss possible future directions for this exciting field.

## Historical introduction to adaptive landscapes

Adaptive landscapes, first introduced in the 1930’s by Sewall Wright [1], are commonly used in evolutionary biology to describe the mapping between genotype and fitness. Rugged landscapes include multiple peaks, or fitness maxima: allele combinations that differ in two or more genes and are superior in terms of fitness to intermediate allele combinations. Natural selection would tend to drive a population towards the nearest adaptive peak, but in many cases this would be a suboptimal solution. How can a population evolve from one adaptive peak towards a higher one, crossing an *adaptive valley*? To answer this question, Wright developed the *shifting balance theory*, in which a population shifts from one adaptive peak to another one via the sequential effects of mutation, random genetic drift, migration, and natural selection [2]. However, Wright’s theory has been criticized for its generalization of the intuitive three-dimensional model to a more realistic high-dimensional genotype space. R.A. Fisher argued in a private correspondence with Wright that the high-dimensional nature of adaptive landscapes means genotypes have many genetic “neighbors”. Thus the probability of any point in the genotype space constituting a local maximum, by not having any genetic neighbors with higher fitness, decreases sharply with the number of dimensions [3, 4].

The conceptual innovation posed by Wright has ignited many theoreticians to try and characterize the potential topology of adaptive landscapes. In turn, the development of adaptive landscape models led to numerous theoretical analyses of evolution on rugged landscapes under different biological conditions. The theory suggests that properties of the adaptive landscapes are major determinants of the progress and outcomes of evolutionary processes. The advent of high-throughput experiments and omics data over the last decade, especially in microbiology, is rapidly changing our understanding of adaptive landscapes. Experimenters are now generating empirical adaptive landscapes by measuring the adaptive values of large sets of allele combinations. These new empirical findings are leading to novel research on the properties of adaptive landscapes and evolution on realistic landscapes. Moreover, the concept and research of adaptive landscapes has transcended biology and is broadly used in computational problems, since finding the highest peak in an adaptive landscape is analogous to finding a global optimum in a multidimensional optimization problem. One example is a spin glass model, which represent the spins of atoms in a magnet. Finding the lowest energy configuration of spins is analogous to finding a global optimum in a rugged landscape, as neighboring spins interact and affect each other and the energy of the system in complex ways [5, 6]. Other examples, from computer science, are *evolutionary algorithms*, which use evolutionary processes such as selection, mutation, and recombination to “evolve” populations of optimal solutions to various problems [7]. Advances in theoretical characterizations of adaptive landscapes and exploration of conditions allowing their efficient traversing can lead to advances in *evolutionary algorithms* [7, 8].

We start this Key Issues Review by introducing several models of adaptive landscapes. We then describe some of the theoretical studies of evolution on rugged landscapes. Next, we review experimental studies focused on measuring empirical adaptive landscapes, and discuss how such empirical studies affect the theoretical literature. Finally, we suggest how future research in this field can integrate empirical and theoretical methods to tackle important issues across science.

## Models of adaptive landscapes

Consider a genotype with *N* loci, or genes, denoted by *g* = (*g*_1_, …, *g*_N_), where *g_i_* represents the identity of the allele at locus *i*. For any *g*, we define a fitness mapping *ω*: ℝ ^*N*^→ ℝ^+^, so that *ω* (*g*) gives the fitness of genotype *g*. The mapping *ω* between genotype and fitness can be modeled in many ways, which can change the possible interactions between loci, *i.e.* generate different epistasis patterns. Excluding the trivial case (*ω* is constant), the simplest definition of *ω* is the ***additive model***: 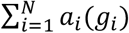, where *a_i_*: ℝ → ℝ is the additive fitness contribution of the allele *g_i_* at locus *i*. A single allele change at locus *j* in the additive model will change the fitness of a mutant *g^*^* only by the marginal difference in fitness conferred by allele 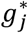:

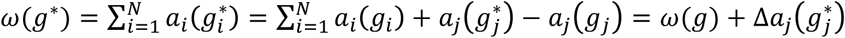

Thus, due to the lack of interactions between loci, an adaptive landscape generated by the additive model is always single-peaked (*i.e.* single maximum). The contrasting scenario is given by a mapping *ω* which assigns a different fitness value for each genotype. This mapping is referred to as the ***House of Cards* (HoC) model** [9] (the name derives from the notion that every mutation can destroy the “biochemical ‘house of cards’ built up by evolution” [10]). The HoC model can be represented by:

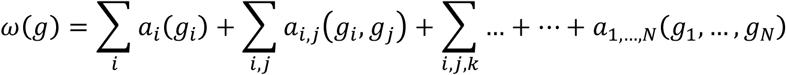

Under the HoC model, each allele combination potentially has a different fitness value, determined by the 2^*N*^ − 1 fitness functions *a*. These are analogous to interactions between variables in a statistical linear model, including main effects (*a_i_*), second-order interactions between genes (*a_ij_*), and so on up to the *N*^th^ order interaction [11]. Thus, the HoC model implies absolute independence between the fitness values of any two genotypes.

A different model that can be extended to include more complex landscapes than additive ones, but still retain a global fitness maximum, is the ***Rough Mt. Fuji* (RMF)** model [12, 13]. RMF is based on an hybrid of the Mt. Fuji model (an additive model containing a single peak, like the famous Japanese mountain [14]) and a HoC model with random coefficients. The fitness of a general genotype *g* will is defined by:

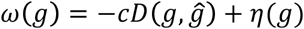

Where *ĝ* is the fittest genotype with respect to the additive model, *D* is a distance metric on genotypes (*e.g.* Hamming distance), *c* > 0 is a scaling constant, and *η*(*g*) is a vector of 2^*N*^ independent identically distributed random variables, assigning fitness values to all gene interactions. If the standard deviation of *η* is much bigger than *c* (σ(*η*) ≫ *c*) then the landscape is governed by the random interactions and the model is reduced to the HoC model. On the other hand, if *c* is much larger than *σ*(*η*), then the model is effectively an additive model. Figure 1 illustrates such a 2-dimensional landscape. The fittest genotype before noise addition (*g ̂*) is found at the center of the horizontal plane (note that noise can shift the location of this genotype). Without noise (*σ*(*η*) = 0) the landscape is completely smooth, taking the shape of the Euclidean distance function, or more generally, the additive model (Figure 1A). Moderate noise (*c* = 1, *σ* (*η*)= 0.1) retains a well-defined peak, but introduces some plateaus and small local peaks (Figure 1B). With substantial noise (*c* = 1, *σ* (*η*)= 0.5) the landscape is extremely rugged with several high local peaks (Figure 1C).

**Figure 1.**
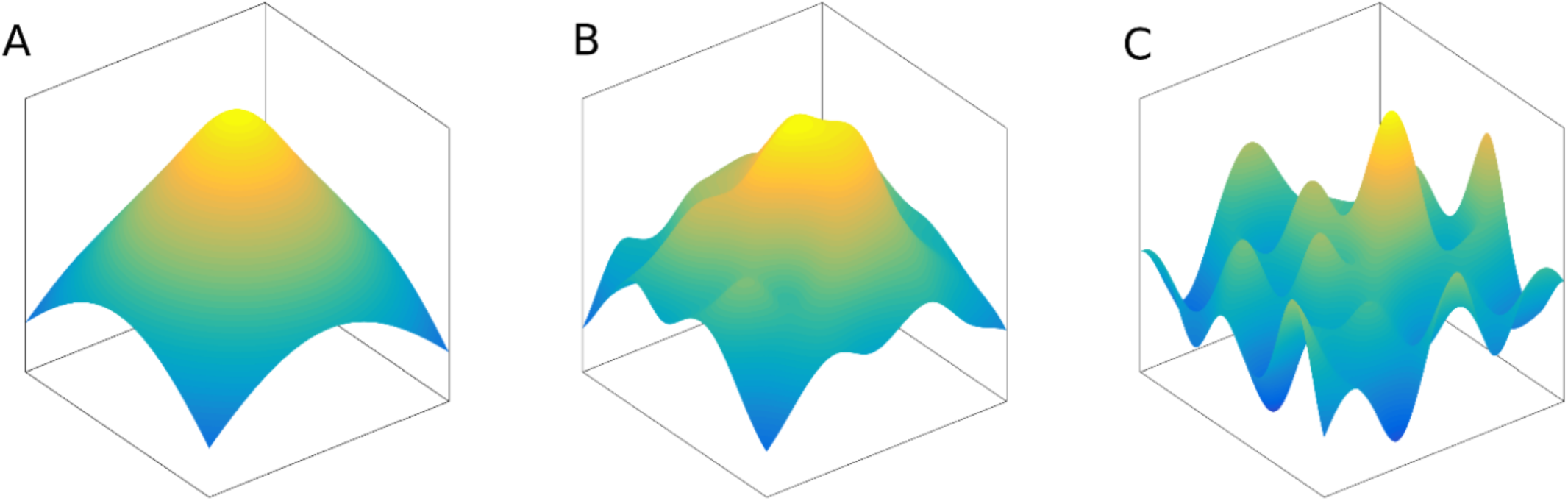
The Rough Mt. Fuji model. The x- and y-axes represent genotypes, whereas the z-axis and color represent fitness. The fittest genotype with respect to the additive model (*g ̂*) is found at the center of the horizontal plane. **(A)** No noise (*σ* (*η*) = 0) yields the additive model. **(B)** Moderate noise (*c* = 1, *σ* (*η*) = 0.1) keeps a similar topology to the noiseless landscape, albeit less smooth. **(C)** Extreme noise (*c* = 1, *σ* (*η*) = 0.5) creates a rugged landscape with several local peaks and shifts the location of the global maximum.

From a biological perspective, these models might not capture essential characteristics of biological systems in which only sets of genes have direct interactions. The ***NK model*** was specifically designed for this purpose. Each of the N genes in this model interacts with K other genes. Consequentially, the additive and HoC models are the limiting cases for *K* = 0 and *K* = *N* − 1, respectively. Unfortunately, intermediate scenarios of the NK model, where 0 < *K* < *N* − 1, are very complex and it is difficult to estimate various features of the landscape, such as the expected number of adaptive peaks and their height [15–17]. We demonstrate the landscapes that emerge from the NK model, and their unpredictable nature in Figure 2. We model a genotype g by a bit-string (comprising either zeros or ones in each locus) of length N=5 (for example, 01100 is a possible genotype). The fitness of each genotype is the average of the fitness contributions of its alleles: 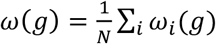. The fitness contribution of an allele depends on the identify of the allele and the identity of *K* other alleles; for example, with *K*=2, we might have *ω*_2_(*g*)= *f* (*g*_0_,*g*_2_,*g*_4_). For this illustration, we randomly chose the K loci that affect each locus, and drawn uniformly distributed random numbers between 0 and 1 for the fitness contribution of each allele, 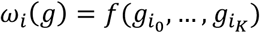 Figure 2 shows such landscapes with *K* between 0 and 4. As expected, *K* tunes the ruggedness of the landscapes: with *K*=0 the landscape is additive and smooth, and higher *K* values are associated with landscapes that are more rugged, as can be seen, for example, by the increasing number of local optima. The source code and step-by-step instructions for creating an adaptive landscape based on the *NK model*, as well as recreating Figure 2, are provided in the online supplementary material (https://github.com/yoavram/UnderTheRug).

**Figure 2.**
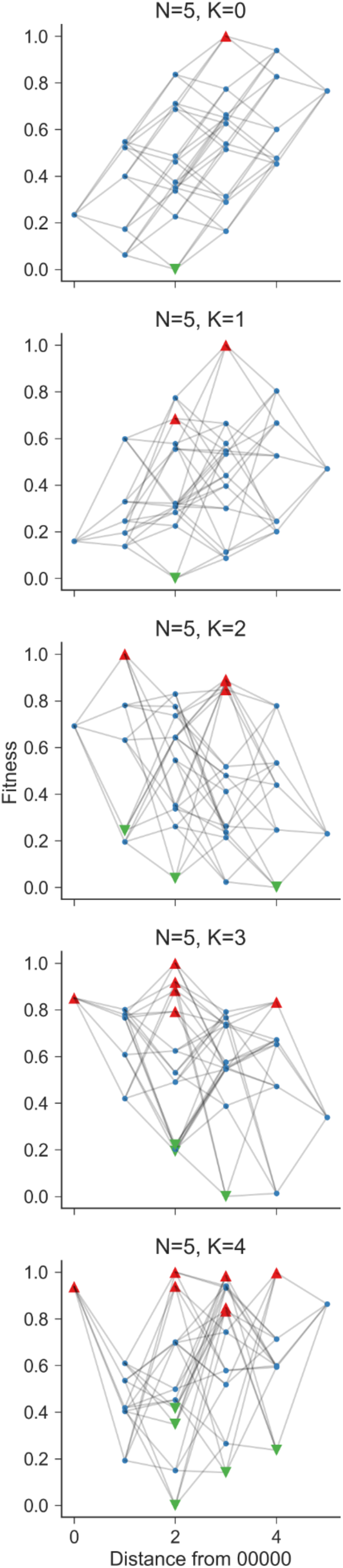
Tunable ruggedness in NK model adaptive landscapes. The figure shows landscapes generated by an NK model with *N*=5 and *K* between 0 and 4, and fitness values of allele combinations assigned randomly (as explained in the main text). The y-axis shows the fitness; the x-axis shows the Hamming distance from genotype 00000 (arbitrary reference point); Markers show the adaptive peaks (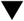), valleys (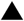), and other genotypes (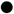); Lines connect genotypes that differ in a single locus. See online supplementary material (https://github.com/yoavram/UnderTheRug) for source code.

The above models attempt to expose the mathematics underlying the topology of rugged adaptive landscapes. However, is ruggedness truly a feature of realistic adaptive landscapes? The notion that adaptive valleys and peaks are sparse in high-dimensional landscapes (*N*>>1) was known from early comments made by Fisher [3], and has been considered an overlooked factor that might drive evolutionary dynamics on seemingly rugged adaptive landscapes [18].

One of the theories implementing this notion of sparsity is the ***holey adaptive landscape theory***, formulated by Sergey Gavrilets [19, 20], where only two fitness values exist: the fitness of each genotype can be either 1 (viable) or 0 (non-viable), and “ridges” can connect highly fit genotypes [21]. The holey adaptive landscape theory relies on *percolation theory*, which describes the connectedness of discrete multi-dimensional spaces: consider a multi-dimensional lattice in which each point is independently assigned 1 with probability *P* and 0 with probability 1 – *P*. Percolation theory predicts that there exists a critical parameter value, *P_c_* (the value of which depends on the number of dimensions), above which there is likely to exist a large connected component of neighboring 1’s on the lattice [22]. A three-dimensional example of percolation theory is given in Figure 3. We simulated 30×30×30 lattices and plotted the average relative size of the largest connected component in them, as a function of *P* (Figure 3A). A steep increase in the size of the largest component is seen once *P* is larger than the percolation threshold (dashed vertical line, Figure 3A), which is ≈ 0.31 in this setting [23]. Cross sections of two simulations show the substantial increase in connectedness when increasing *P* from below to above the percolation threshold (Figure 3B and 3C, respectively).The source code and instructions for recreating Figure 3, are provided in the online supplementary material (https://github.com/yoavram/UnderTheRug).

**Figure 3.**
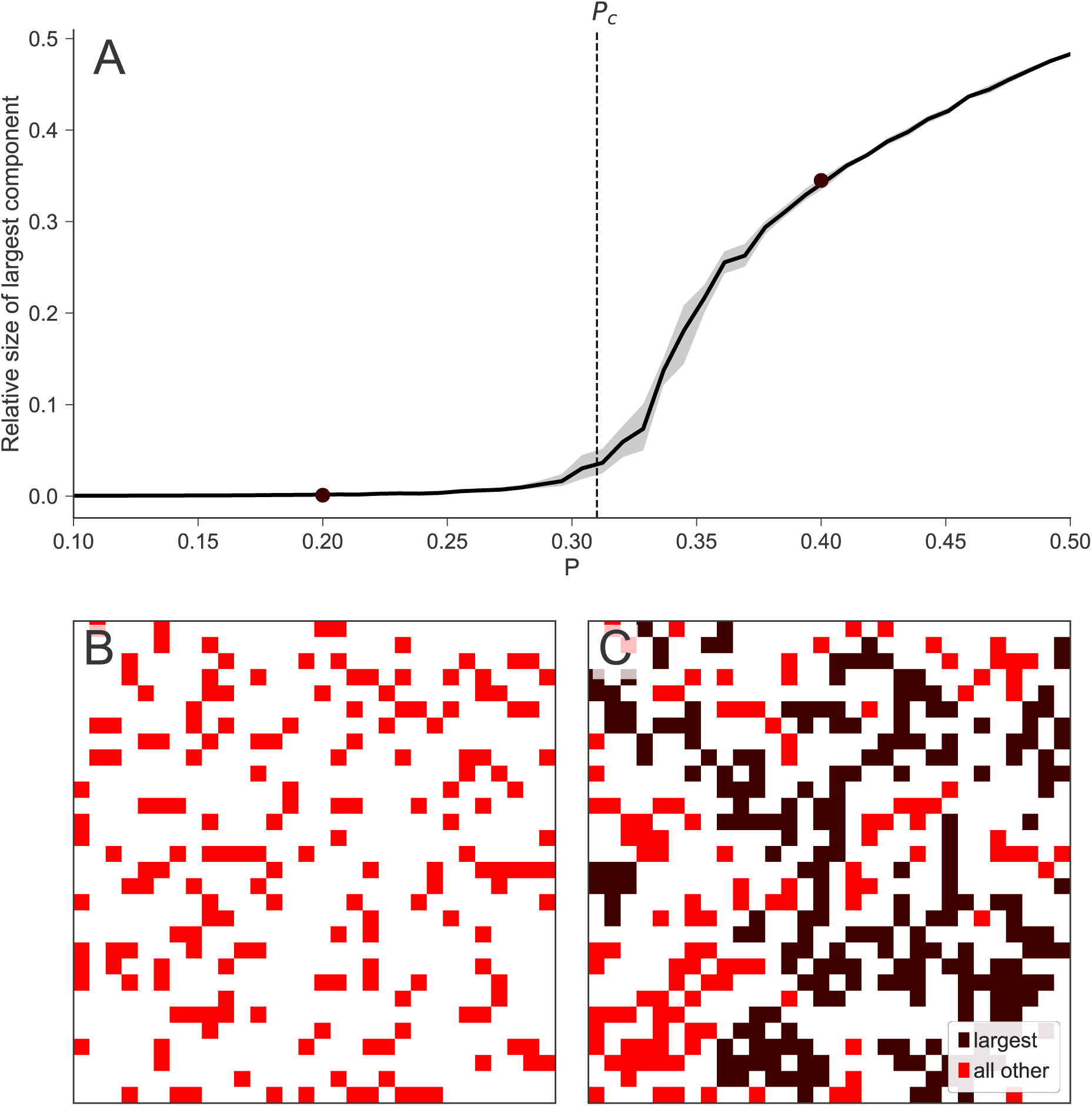
Connectedness of a holey landscape. (A)The size of the largest connected component in a random 30×30×30 lattice relative to the lattice as a function of *P*, the probability that a lattice cell is 1 rather than 0. The critical probability *P_c_* = 0.31, below which the largest component is very small, is represented by the dashed line. The solid line shows average value for 10 random lattices, narrow shaded area shows a bootstrapped 99% confidence interval. **(B)** a cross section of a random 30×30×30 lattice with P=0.2. **(C)** a cross section of a random 30×30×30 lattice with P=0.4. Dark color shows the largest component and light color shows all other viable components. The largest component in C constitutes 33% of the lattice, whereas the largest component in B, with P=0.2, consists of only 0.2% of the lattice and therefore isn’t seen in the figure. See online supplementary material (https://github.com/yoavram/UnderTheRug) for source code.

Gavrilets has drawn the analogy from percolation to adaptive landscapes. When we consider the lattice to be the genotype space and the binary outcomes as high and low fitness values, the landscape will look like a slice of Swiss cheese riddled with holes, hence *holey*. If the probability for genotypes with high fitness exceeds *P_c_*, we can expect evolutionary trajectories between almost any pair of high fitness genotypes to circumvent the “holes” - the low fitness genotypes. Later, Gavrilets and others have extended the theory for continuous and correlated fitness landscapes [20].

The *holey adaptive landscapes theory* explains how most fit genotypes could appear and reach high frequencies due to mutation and natural selection under several conditions: First, the observed values of *P* should be higher than the critical value, *P_c_*, a condition that is difficult to test and that might not be satisfied in many scenarios. Second, if ridges connecting high fitness genotypes are “narrow” enough, evolutionary trajectories between these genotypes can be very rare, and additional evolutionary processes are likely needed to explain adaptation. This relates to the more general problem of the characterization of the evolutionary accessibility of adaptive landscapes (*i.e.* existence paths in genotype space in which each mutation increases the fitness) [24–27]. So the question remains – how do populations reach evolutionarily inaccessible adaptive peaks?

## Models of evolution on adaptive landscapes

The development of adaptive landscape models inevitably raised the question of how populations cross adaptive valleys: if different mutations are separately deleterious but jointly advantageous, how can a population evolve from one adaptive allele combination, or peak, to a fitter one, crossing a less fit valley? A *peak shift* process can be described as consisting of two stages: the emergence of a genotype on the higher peak, and the spread and fixation of this genotype in the population [28]. Facilitating each of these stages usually requires conflicting evolutionary conditions.

Intermediate genotypes are usually disfavored by natural selection, because these genotypes are deleterious or neutral compared to the common genotypes. Since the emergence of a genotype on a higher adaptive peak is usually a result of mutation or recombination in intermediate genotypes, large numbers of intermediate genotypes will hasten the emergence of the adapted genotype. Therefore, strong selection will tend to postpone the appearance of the adapted genotype. However, high mutation rates (*i.e.*, the population-wide rates of mutation) may allow even few intermediate genotypes to produce the adaptive change in a process called *stochastic tunneling* [29–31].

In contrast, weak selection can help the emergence of the adaptive genotype as it allows a larger number of intermediate genotypes to survive; however, with weak selection, the advantage of the adaptive genotype is small and it is therefore more likely to go extinct due to random drift. Due to these conflicting conditions for each stage, the expected time for a peak shift can be very long, especially in small populations where the adaptive advantage of the higher peak is small, or in cases where the adaptive disadvantage of intermediate genotypes is large [32–34].

Wright tried to solve this problem, which he was also responsible for posing, using the *shifting balance theory of evolution* (SBT) in 1931 [1]. Wright proposed that in a population divided to small subpopulations, each subpopulation is subject to random genetic drift, and may therefore stochastically descend from an adaptive peak to a valley. When the population is found in the valley, mutation or recombination can produce new, adaptive genotypes which will be favored by selection, effectively driving the subpopulation up to a nearby, possibly higher, adaptive peak. Thereupon the adapted subpopulation can send immigrants to “takeover” other subpopulations. The first three phases (drift, mutation, and selection) are illustrated in Figure 4, where the horizontal plane represents genotypes, and the height and color represent fitness. We show how a small population from a local peak can produce maladapted mutants via mutation (Figure 4A); these mutants increase in frequency by drift, and create the fittest genotype, found on the global maximum, by recombination or mutation (Figure 4B); the fittest genotype can reproduce and increase in frequency (Figure 4C), outcompeting other genotypes due to its superior fitness (Figure 4D). The time scales of these processes are represented by the clocks on Figure 4. Whereas selection driven processes are fast (Figure 4B-D), mutation and drift driven processes are typically much slower (Figure 4A) [32–34].

**Figure 4.**
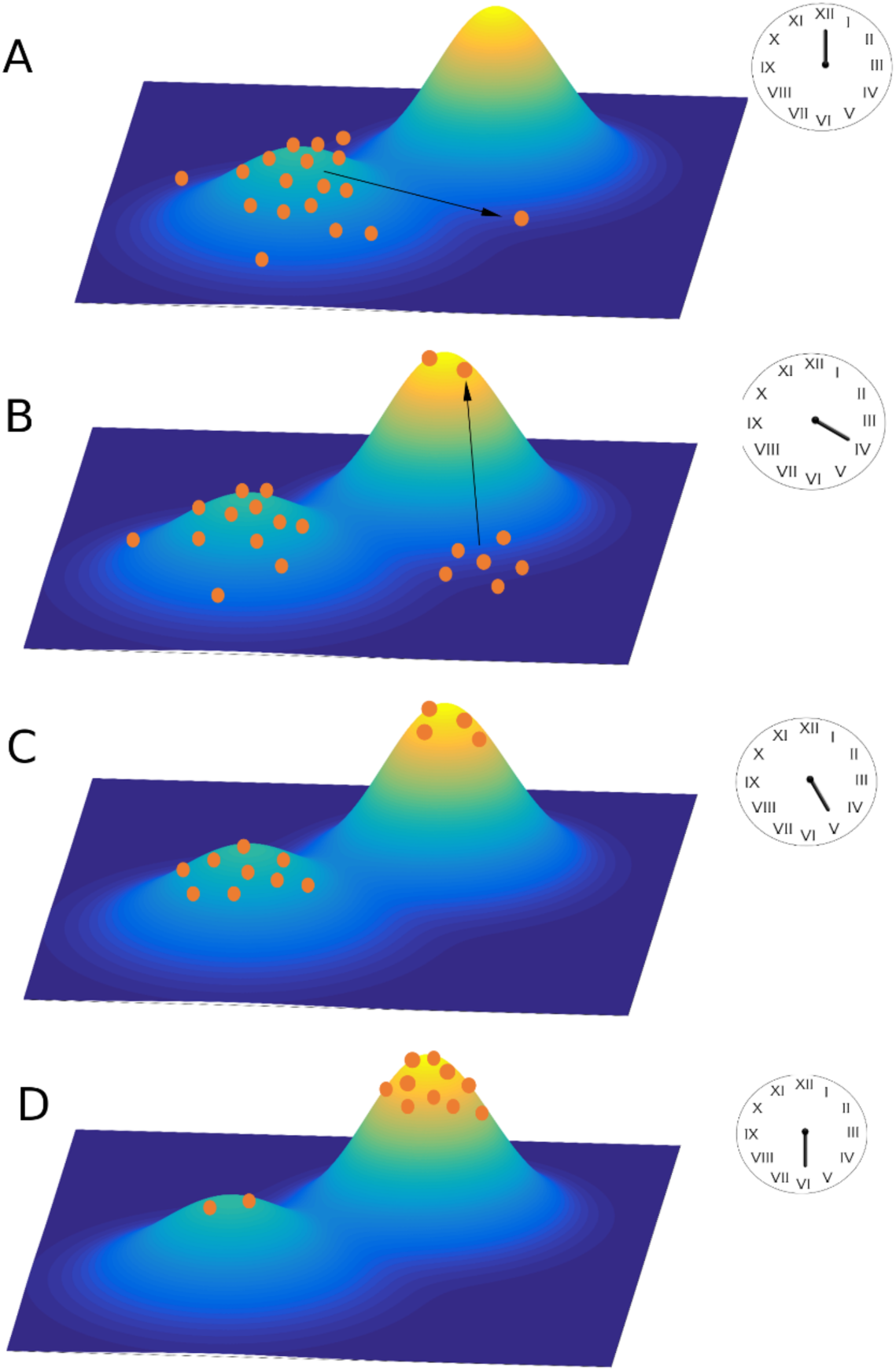
Illustration of peak shift dynamics. Genotypes are given by the x and y axes, the fitness of each genotype is represented by height and color, and individuals are marked by orange circles. Clocks loosely represent the duration of evolutionary processes. **(A)** The initial population is distributed around a local adaptive peak, and maladapted mutants appear by mutation (arrows) and can increase in frequency by random genetic drift. **(B)** Maladapted individuals at the base of the global adaptive peak can produce adapted individuals by mutation and/or recombination (arrows). **(C)** Adapted genotypes fix and proliferate due to natural selection and **(D)** eventually take over the population.

We demonstrate such a peak shift process using a relatively simple landscape with epistasis between just one pair of loci: allele combinations *ab, aB, Ab,* and *AB* contribute *1, 1-s, 1-s*, and *1+sH* to the fitness; 25 additional background loci have multiplicative effects, where each deleterious allele reduces fitness by 1-s. Therefore, *ab* and *AB* are local and global adaptive peaks, respectively. We simulate peak shifts by initiating a population on the lower peak *ab* and letting it evolve towards the higher peak via mutation, drift and selection (Figure 5A). The peak shift process is divided into three distinct time periods: (i) the appearance of a double mutant - an individual with *AB* genotype (Figure 5A blue) - driven by drift and mutation, and therefore taking a long time (Figure 5C) [35, 36]; (ii) the possible extinction of the double mutant while it is rare (Figure 5B) due to random genetic drift - this can be usually modeled as a branching process [37]; and, if the double mutant avoided extinction (iii) the time to fixation of the double mutant due to natural selection, which is much shorter than the appearance time (Figure 5D) [38]. The data used to create Figure 5C-D was taken from [39] (see source code and details in the online supplementary material (https://github.com/yoavram/UnderTheRug).

**Figure 5.**
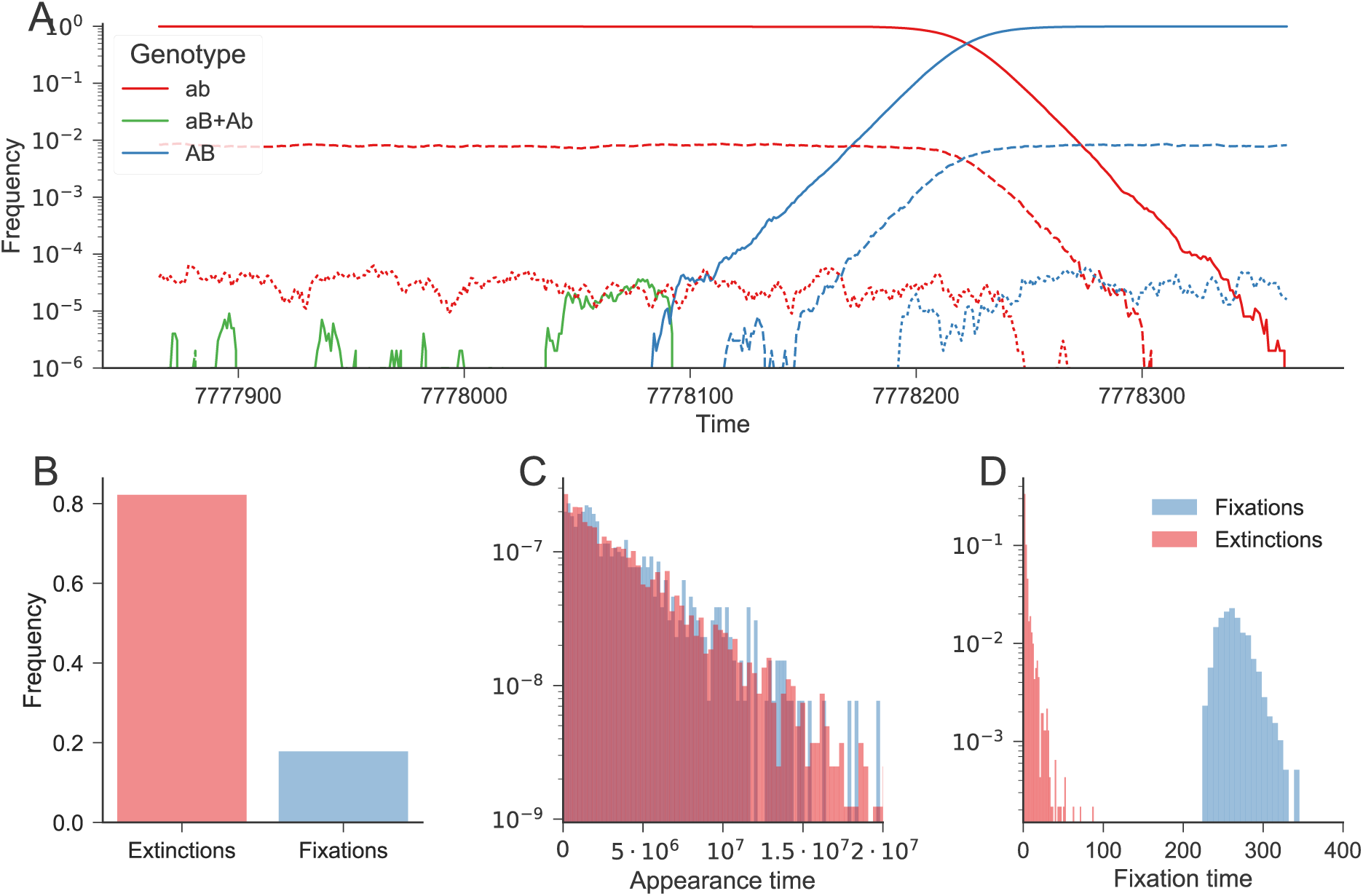
Peak shift simulations on a rugged adaptive landscape. Populations started with genotype *ab* and fitness *1* and evolved to genotype *AB* with fitness *1+sH* by crossing the adaptive valley of genotypes *aB* and *Ab* with fitness *1-s*. **(A)** Frequency of different genotypes during the final 500 generations of an adaptive peak shift. Colors denote the genotype at the two loci of interest (*ab* in red; *aB* and *Ab* together in green; *AB* in blue); line styles denote the number of deleterious mutations in the background (solid: zero; dashed: one; dotted: two). The frequency of the intermediate genotypes (green) increases around generation 7778050 due to drift, which leads to the appearance via mutation of the double mutant (blue) around generation 7778090. The double mutant avoid extinction and then increases in frequency due to selection, eventually becoming the dominant genotype (around generation 7778220). **(B)** Frequency of fixations (blue) and extinctions (red) of the double mutant after first appearing. **(C)** The distribution of the time until the double mutant first appears in the population in simulations ending in fixation (blue) or extinction (red) is roughly exponential (note the log scale). **(D)** The distribution of the time until fixation (blue) or extinction (red) of the double mutant after it appears. Parameters: 10^6^ individuals; selection coefficients *s*=0.05, *sH*=0.1; mutation rate at focal loci 8 ∙ 10^−8^ mutations per base-pair per generation; mutation rate at background loci 4 ∙ 10^−4^. In total, 3,246 samples were simulated, with 578 fixations and 2,668 extinctions. See online supplementary material (https://github.com/yoavram/UnderTheRug) for simulation and analysis code.

Recently, Bitbol and Schwab generalized the SBT. Their analysis showed that subdivision of a population into smaller subpopulations, connected by migration, is enough to accelerate the peak shift process under certain parameter ranges [40]. In contrast to SBT, they do not require that the volume of migration from subpopulations will depend on their inhabitants’ fitness. Despite this generalization of SBT, it is only relevant under specific biological constraints (*e.g.* subdivided population, sequential appearance of mutants, etc.), which might not apply to a vast range of biological scenarios. Some critics even hold that peak shifts under the SBT framework have not played a significant role in evolution [41, 42], although evidence to the contrary also exists [43, 44]. As a reaction to SBT’s inability to account for peak shifts under diverse circumstances, a plethora of theoretical studies have offered various environmental conditions, biological mechanisms, and evolutionary forces that could facilitate adaptation on rugged adaptive landscapes.

Michael Whitlock has suggested a peak shift process complementary to Wright’s SBT: *variance induced phenotypic shift* (VIPS) [45]. In the first stage of SBT, small populations are assumed to have decreased genetic variance in comparison to larger populations (which effectively increases drift and allows intermediate maladapted genotypes to appear and survive). However, as Whitlock noticed, the amount of variance in small populations is itself a random variable, which could have high variance. Therefore, some of the populations are bound to have higher variance then the expected value Wright assumed. Whitlock showed (based on a model and an observation made in [46]) that the mean adaptive landscapes of populations with high variance could be substantially different from the adaptive landscapes of single individuals. Specifically, he proved that increased variance could lead the mean fitness of a population to be a unimodal distribution around the individual’s fitness maximum. Thus, selection could deterministically lead the population across the fitness valley. Notably, Whitlock has also demonstrated that rugged individual fitness maps can be translated into unimodal mean adaptive landscapes even with low variance, if a small environmental perturbation changes the distribution of the population [47].

Other researchers have tried to characterize possible biological scenarios reconciling the seemingly contrasting conditions needed for a peak shift. Wright conceived that changes in environmental conditions could remodel the adaptive landscape in ways that may aid a peak shift [2, 48]. More specifically, Hadany has showed that peak shifts can be much more common in subpopulations that experience different intensities of selection and are interconnected by migration [49]: The adaptive genotype could emerge in the population experiencing weak selection, due to large number of intermediate genotypes, then migrate to the second population, and spread there due to the harsher selection regime.

Ram and Hadany suggested that a mutator allele that increases the mutation rate specifically in low-fitness individuals, found below the adaptive peaks, can significantly decrease the time for an adaptive peak shift. Furthermore, such an allele will not jeopardize the fitness of the population after adaptation is achieved, which is the main disadvantage of standard mutator alleles [35]. Similarly, Peak shifts are facilitated when there is a negative association between fitness and the generation of variation: recombination [50], dispersal [51], or viral coinfection [52].

Cooperative behavior has also been recently examined as a factor that might accelerate crossing of adaptive valleys, in contrast to previous research, focusing mostly on biological mechanisms and environmental constraints. In a *division of labor* type of cooperation, wherein individuals contribute unique products necessary for the population survival, a mutant capable of producing several products was found to rapidly arise as a response to “cheaters” that do not produce any goods [53, 54]. Another study used a *public goods* cooperative model, where individuals contribute a portion of their fitness and redistribute it among all group members. Such cooperative behavior effectively “flattens” the adaptive landscape by redistributing fitness from the fit to the unfit, thereby reducing selection against intermediate genotypes. This accelerates the emergence of genotypes at the adaptive peak, although it may also hinder their spread [55].

Other notable studies applied and extended the concept of adaptive landscapes without directly referring to peak shifts: Bak et al. [56] used an extension of the *NK* model to study the coevolution of community of mutually dependent species. This extension is called the *NKC* model [57], as the fitness of a species is a function of *N* of its genes and *C* genes of other species. They showed that the model can result in two distinct scenarios, depending on the relationship between *C* and *N*. In the first scenario, every species finds a local adaptive peak and evolution essentially stops. In the second scenario, only some species are at a local peak. Other species are evolving towards a peak, but in doing so, they change the landscape, effectively pushing already adapted species off their peaks. In this scenario, therefore, evolution can proceed, perhaps indefinitely.

Kryazhimskiy et al. [58] developed a general theory to analyze adaptation on different types of adaptive landscapes, including the HoC landscape and smooth landscapes such as the *Stairway to Heaven* landscape, so called because the distribution of the fitness effects of new mutations is the same for all genotypes, allowing adaptation to proceed indefinitely. They found that adaptation dynamics can be characterized by the expected fixation probability and fitness increment of new mutations. They suggest a classification of adaptive landscapes per the emerging adaptive dynamics and demonstrate how landscapes can then be inferred from fitness data.

Using simulations with in the *Avida digital evolution* platform [59], Clune et al. found that on smooth landscapes natural selection can optimize mutation rates with respect to fitness, but not on rugged landscapes with “wide” adaptive valleys. The mutation rate must balance between the ability to adapt, increased by the rate of beneficial mutation, and the ability to remain adapted, decreased by the rate of deleterious mutation [60]. The authors show that the reduced likelihood of beneficial mutations on rugged landscapes tips this balance towards elimination of deleterious mutations, thus limiting adaptation in the long term.

## Empirical adaptive landscapes

Observations of epistatic gene interactions in a limited number of loci and alleles have been described as early as the beginning of the 20^th^ century [61]. However, the technological leaps in molecular biology methods towards the end of the 20^th^ century and the beginning of the 21^st^ century, most notably the genomic revolution [62, 63], have significantly changed the research of adaptive landscapes. For the first time, large empirical adaptive landscapes could be comprehensively characterized and studied.

One of the earliest studies characterizing adaptive landscapes directly from known mutations (as opposed to indirect measures of mutations, relying on supposed relationships between a number of mutations and fitness, inbreeding coefficients, etc. [64–67]) was performed by de Visser et al. in 1997 [68]. They studied synergistic epistasis between mutations in the fungus *Aspergillus niger* in an attempt to test the claim that sexual reproduction is advantageous because it breaks combinations of deleterious mutations [69, 70]. Using a set of mutations in eight genes, conferring drug resistance, auxotrophy, and spore color, they measured the fitness of the fungus with all 256 allele combinations. Although not all mutants were observed (only 186 of 256 possible allele combinations were viable), the sampling was sufficient to create an adaptive landscape. Interestingly, the fitness was found to be overall log-additive in the number of deleterious mutations, but this was the result of synergistic and antagonistic effects of different mutations cancelling each other. Generalizing conclusions from such results is precarious. The lack of epistasis in a small set of loci is not necessarily indicative of the interactions between other loci. Moreover, sets of mutations within the same gene may have different properties compared to sets of mutations across the genome. Furthermore, mutations in this study were only defined by their phenotype. Although changes in phenotype are consistent with discrepancies from wild type alleles, different mutations leading to the same phenotype could have had different effects on mutants’ fitness. This confounding effect could not have been regarded without genetically characterizing each mutant.

Many other early studies [64–67] characterized empirical adaptive landscapes but were constrained by the number of mutants they could examine and the lack of certainty regarding the mutants’ genotypes. Such caveats emphasize the difficulty of constructing general and reliable empirical adaptive landscapes, especially before the era of high-throughput genetic data.

One way of reducing the complexity of estimating adaptive landscapes, but retaining high dimensionality, is focusing on single macromolecules rather than entire organisms. Fitness can then be defined as a molecule’s performance of a certain function, serving as a proxy for the fitness of the organism that produces this molecule. This method has been applied on various protein landscapes [71- 78] since many variations of them can be produced and their fitness measured in bulk. For example, Wu et al. [72] have characterized the adaptive landscape of an immunoglobulin-binding protein expressed in Streptococcal bacteria. They constructed all 20 amino acid variants in four sites of the protein (leading to 20^4^=160,000 variants) and measured the stability of the immunoglobulin’s and its affinity to a receptor protein. When they restricted the amino acid variants to a small subset, or when they diminished the number of loci examined, sign-epistasis obstructed most increasing evolutionary trajectories. However, relaxing the constraints smoothed the adaptive landscapes and allowed for easier adaptation through the dimensions added.

RNA molecules are also a popular alternative to whole organisms when measuring adaptive landscapes [79–84]. Moreover, they are even smaller and simpler to handle in bulk quantities than proteins. Jiménez et al. have estimated the adaptive landscape of a short RNA molecule (24 nucleotides) by creating all possible sequence variations [81]. Fitness was defined by *in vitro* selection of sequences best sequestering GTP, the rationale being that GTP served as an energetic currency in the ancient stages of evolution, when RNA was the main form of genetic material available. The measured adaptive landscape was extremely sparse, with the frequency of peaks being approximately 10^-13^. This low frequency falls below the percolation threshold *P_c_* and, in accordance to Gavrilets’ theory (see above), peaks were indeed extremely isolated, and only a subset of the peaks were connected by just a few evolutionary trajectories. The colossal number of variants (4HK ≈ 10%K) covered by Jiménez et al. demonstrates the problem of measuring empirical adaptive landscapes, even for small molecules. Although they have covered all possible variations, the space itself was low-dimensional and consisted of only 24 nucleotides. In another study, Aguilar-Rodríguez et al. measured transcription factor binding affinity of all possible combinations of an 8 nucleotide DNA sequence across many eukaryotic species, from yeast to mice [85]. The study produced over a thousand different adaptive landscapes, but retained the same caveat as the previously mentioned studies: it covered the adaptive landscape of only a small DNA fragment relative to the entire genome.

This raises the question of whether a sample of the molecule space can have substantial predictive power: could we extrapolate the entire molecule space from the effects of mutations sampled in its subspace? A common method is applying regression models [74, 86, 87] to the sampled subspaces, often with regularization to reduce over-fitting [88], to try to extrapolate to higher dimensions.

However, the results of such models might be biased [89]. Plessis et al. have tried to overcome this problem by applying different sampling methods of the available space [90]. They have found that in their dataset, none of the examined methods could accurately estimate interactions of order >2. Furthermore, they found that (i) completely random sampling of the available subspace around a sequence of interest is usually inefficient; (ii) exhaustive sampling of only a subset of dimensions allows a good prediction of the fitness of genotypes close to the sampled region, but poorly predicts fitness of farther genotypes; (iii) random sampling close to the sequence of interest is slightly worse at predicting the fitness of close genotypes, but can extrapolate more easily to farther genotypes than exhaustive sampling; and (iv) sampling genotypes from a population that has undergone selective pressure and contains mostly high fitness genotypes seemed to incorporate the advantages of all other methods of sampling and produce accurate predictions. Despite these remarkable results, the extent of their application to other adaptive landscapes is unknown and would have to be studied.

Of course, even accurate estimates of functional differences between different alleles of a single molecule (*e.g*, affinity of a protein receptor) are only proxies for organismal fitness, which depends on many other molecules and genes, some of which can compensate or mask the differences between said alleles. Indeed, some studies actually use organismal fitness rather than *in vitro* molecule function [82, 91, 92], but measuring organismal fitness is likely to increase measurement noise and introduce experimental bias. The choice of the measured trait and experimental setup, therefore, inherently entails a trade-off between resolution, accuracy, and robustness.

Another approach for estimating adaptive landscapes is to focus on mutations in a single gene or macromolecule, but to measure the effect of mutations on aspects of organismal fitness *in vivo* rather than use the *in vitro* function of the macromolecule as a proxy. Focusing on the bacterial TEM gene that confers resistance to β-lactam antibiotics, Weinreich et al. [76] utilized 5 mutations in a mutant allele that increases resistance to the antibiotic cefotaxime by 100,000-fold to create 2^5^=32 allele combinations. The researchers estimated the resistance provided by the different allele combinations by measuring the minimal concentration of the drug that inhibits growth of bacterial colonies that incorporate each specific allele. Using this approach, they could show that sign epistasis is prevalent in TEM, as the effect of many mutations could be either positive or negative, depending on the identity of the other alleles. Due to the prevalence of sign epistasis, only 18 evolutionary trajectories (i.e. alternative orderings of the 5 mutations), out of a total of *5!*=120, consisted of strictly increasing resistance. Furthermore, applying classical population genetics – that the fixation probability of a mutant genotype is proportional to its advantage over the resident population – they concluded that two of these trajectories are much more likely than the other 16 (**Error! Reference source not found.**6). This suggests that the process of adaptation to cefotaxime, in terms of the order of mutations that are fixed in the adapting populations, is highly predictable. However, a recent experimental study showed that when bacteria were first selected by intermediate and then by harsh antibiotic selection, they eventually reached different, and even higher, peaks than those reached in the TEM landscape imposed by harsh selection alone [93]. These results confirm that environmental change can affect peak shifts substantially, and that the evolutionary predictability is conditioned on the environmental context.

A similar approach was taken by de Vos et al. [94], who focused on the *lac* operon, responsible for lactose metabolism when and only when lactose is the major carbon source in the media. They measured the expression of the *lac* proteins in 64 *lac* repressor and operator mutants. The resulting rugged empirical landscape consists of 720 alternative evolutionary trajectories. The authors found that if the environment is stable, the population is stuck in suboptimal local peaks. But if the environment fluctuates between two conditions – with and without lactose – adaptation can proceed. They demonstrated that specific genotypes that constitute “adaptive valleys” in one environment become “peaks” in the other environment; thus, environmental fluctuations allow selection to drive the adaptive process without the long waiting times associated with valley crossing (as previously suggested by Wright [2] and Whitlock [47]).

## Combining empirical landscapes with theory

For many years, evolutionary biologists could only reason and extrapolate about the topology of adaptive landscapes. Therefore, most evolutionary models used generalized and mathematically convenient mappings between genotypes and fitness. However, the increasing availability of empirical landscapes allows to examine their topological properties and to compare them to theoretical landscape models. Two recent review papers have performed meta-analyses of published empirical landscapes’ characteristics [95, 96]. They examined how features such as the number of fitness peaks, the frequency of epistatic interactions and the ‘ruggedness’ of the landscape correlate in empirical data, and how they would be expected to correlate under theoretical landscape models. They found that landscapes’ topologies are substantially more rugged than those expected under an additive model, but less than ones expected under a completely random interaction model (*e.g.* House of Cards). Furthermore, a rough Mt. Fuji model was able to replicate the association between topological characteristics of the observed landscapes well [96], suggesting that it might provide a realistic way of generating landscapes on which evolutionary theory could be studied. Hence, modelers can now either simulate evolution using estimated statistical and topological properties of empirical landscapes to construct generalized realistic landscapes, or directly simulate on specific empirical landscapes.

We demonstrate the latter approach by simulating adaptive evolution on the *A. nigeri* [68] and the TEM [76] landscapes, using the classical Wright-Fisher model [97]. This model is widely used in population genetics because it captures many biological scenarios and is more computationally efficient than other methods, such as individual-based simulations and Moran processes [97]. The Wright-Fisher model follows the frequency of different genotypes over time. Here, we focus on an asexual, haploid population with a constant population size, undergoing natural selection, random genetic drift, and mutation. The effect of natural selection is defined as:

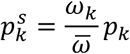
where *p_k_* is the frequency of genotype *k* before selection, *ω_k_* is the fitness of genotype *k*, and *ω̅* =Σ_*j*_ *ω_j_* ⋅ *p_j_* is the population mean fitness. The effect of mutation is modeled by:

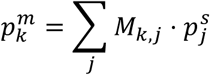

where *M_i,j_* is the mutation rate from genotype *j* to genotype *i* and ∀*j*, Σ_*i*_,*M_i,j_*=1. Finally, the effect of random genetic drift is given by

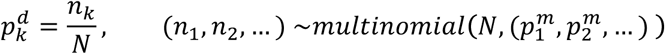

where *N* is the population size, and *n_k_* is the number of individuals of genotype *k* after drift, drawn from a multinomial distribution with *N* trials and probabilities *p^m^*, determined by the frequencies after the effect of selection and mutation. Computationally, selection and mutation can be implemented with matrix multiplication followed by division by *ω̅*, which is the inner product < *ω*, *p* >. Drift can be implemented by drawing from a multinomial random number generator and dividing by the population size (see SI for the source code of a Python implementation).

Empirical adaptive landscapes provide the simulation with the set of genotypes (*k*) and their fitness values (*ω*); however, we still need to determine the mutation rates between these genotypes (*M*). Mutation rates are hard to measure [98] and can vary between species, populations, individuals and even loci on the same genome [99]. In the case of the TEM landscape, we can approximate these using the *E. coli* genomic mutation rate, which has been estimated in numerous studies (see [100–102] for recent estimates), as the genotypes differ by a single amino-acid, although this approximation still neglects some factors. In the case of the *A. nigeri* landscape, we have a greater problem, as we are not aware of the DNA sequences underlying the different alleles, nor do we have estimates for the mutation rate in this organism. That being said, we can still set the mutation rate to some reasonable level and accept the results with a “grain of salt”.

Our simulations started with an isogenic population far from the global peak and proceeded until the population adapted. The results demonstrate several evolutionary phenomena. First, in both landscapes adaptation starts with a big leap forward, but additional fitness increases become smaller and farther apart (Figure 6A,D). Similar dynamics have been observed, for example, in bacteria in the Long Term Evolutionary Experiment [103] and in experimental evolution in fungi [104]. Second, in the smooth and single-peaked TEM landscape, every step along the evolutionary trajectory reached high frequency before the next genotype emerged and increased in frequency (Figure 6B). This is because there are several trajectories along the adaptive landscape that are strictly increasing in fitness (Figure 6C) and natural selection is likely to push the populations along these trajectories. Indeed, each subsequent frequent genotype was just one mutation away from its predecessor. In contrast, in the rugged *A. niger* landscape, in which 6% of the genotypes are adaptive peaks, only four genotypes reach a high frequency (Figure 6E), although adaptation proceeds through seven different genotypes. Moreover, these four genotypes differ from each other in 1, 3, and 3 mutations (Figure 6E, bold lines in Figure 6F). On this rugged landscape the population must cross at least two adaptive valleys (Figure 6F), and the intermediate genotypes at the valleys are unlikely to become very frequent. As with previous simulations, we provide the source code and analysis of the simulations in the online supplementary material (https://github.com/yoavram/UnderTheRug).

Similar approaches to simulations on empirical fitness landscapes have recently been used to suggest that recombination accelerates adaptation in HIV [105], to analyze the relationship between population size, mutation rate, and the predictability of adaptation [106], and to test if sex accelerates adaptation on rugged landscapes [107]. However, simulations on empirical adaptive landscapes are still rare, and we consider this new approach very promising.

**Figure 6.**
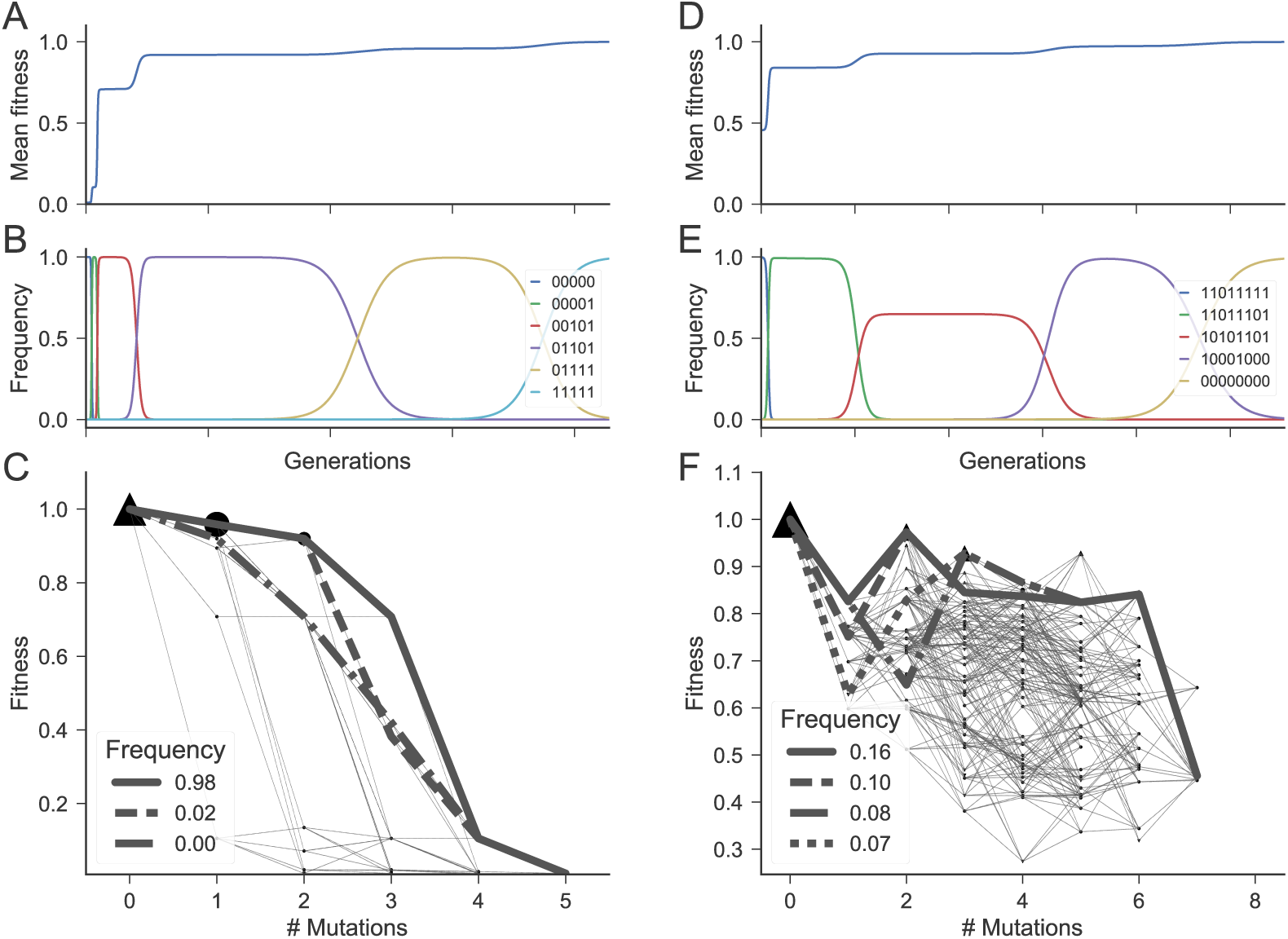
Evolutionary simulations on empirical fitness landscape. Results of Wright-Fisher simulations on two empirical fitness landscapes. **(A-C)** An adaptive landscape consisting of 32 genotypes of the bacteria *E. coli* from a study by Weinreich et al. (2006). Genotypes were constructed from a combination of 5 single mutants in the TEM β-lactamase gene. Simulations started with a population that is 5 mutations away from the global adaptive peak. The mutation rate is 10^-8^ mutations per base pair per generation. **(D-F)** An adaptive landscape consisting of 186 genotypes of the filamentous fungus *A. niger* from a study by de Visser et al. (1997) (the dataset was extracted from Franke et al. (2011)). Genotypes were constructed from a combination of 8 single mutants (including both metabolic and resistance genes). Simulations started 7 mutations away from the global adaptive peak. The mutation rate was 10^−4^ mutations per gene per generation. **(A,D)** The population mean fitness over time in a single simulation. **(B,E)** The frequencies of the most frequent genotypes over time in a single simulation. **(C,F)** Fitness (y-axis) and number of mutations away from the global adaptive peak (x-axis) for all genotypes. Triangles denote fitness maxima (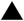); other genotypes are denoted by circles (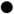); thin dotted lines connect genotypes that differ by a single mutation. The bold lines represent the most common evolutionary trajectories in our simulations, which all started with a single genotype; marker intensities are relative to their frequency in the simulations. Parameters: 10^9^ individuals; 2^15^ generations. See online supplementary material for simulation and analysis source code.

## Future directions

One important application of adaptive landscapes research is in the arms race between pharmacology and pathogens. Drug resistance is an important example: pathogens that quickly adapt to newly developed drugs are a major public health hazard [108–110]. Remarkably, resistance to some antimicrobial drugs was found to correlate with sensitivity to other drugs [111–114], indicating that the underlying adaptive landscape is rugged. This notion can be used to develop treatments regimens that hinder evolution of drug resistance [111, 115, 116]. First steps have been made to design drug regimens based on specific empirical adaptive landscapes related to bacterial pathogens [117–120], malaria [121, 122] and HIV [91]. Further research with additional drugs and increased landscape sampling resolution will be necessary to make this a part of standard practice.

A related concept is that of *evolutionary trapping*: the use of a drug that specifically selects for a particular genotype, ‘setting the trap’, followed by a second drug that specifically selects against that same genotype, thereby ‘springing the trap’. A proof-of-concept has been demonstrated in yeast [123], where radicicol was first applied to drive the gain of an additional chromosome in a population; this chromosome gain increased the sensitivity to hygromycin B, which was then used to exterminate the population. Understanding the complex adaptive landscapes involved in this scenario allowed the authors to predict that escape from the trap is highly unlikely.

Our understanding of cancer as an evolutionary process [124, 125] is to some extent analogous with microbial evolution [126]. An important part of understanding the evolution of malignancy, progression, and metastasis is to measure and model the adaptive landscape on which this evolution occurs.

Eventually, this knowledge could help us design safer and more efficient treatments [127]. As experimental cancer evolution becomes a major research program in both evolutionary biology and in oncology [128, 129], we expect that empirical adaptive landscapes of cancer cells will be published in the next decade.

Symbiotic microbes can also provide an important extension to the adaptive landscape paradigm via the *hologenome* concept, which accounts for the genetic material of both the host and its entire microbial community (also known as the microbiome) [130]. The evolution of the host, per this view, depends on co-evolution with its symbionts, which usually have much shorter generation time and larger population sizes, as well as considerably different ecology and genetics. Novel sequencing and analysis methods now enable researchers to characterize and explore the dynamics of microbiomes over time [131, 132]. Associations between the host immune-related responses and pathways and the microbiome genetic composition have been found, implying that host-symbiont evolution may be tightly linked [132, 133]. Research into host-symbiont adaptive landscapes can significantly affect our understanding of evolution, as well as lead to innovations across various fields, including agriculture, with crop plants [134], fish, and farm animals [135] as hosts; medicine, with human hosts [136, 137]; and conservation, with wildlife hosts [138]. Theoretical models of community adaptive landscapes have already been developed [56], but there is much left to be done to integrate these models with empirical data.

Adaptive landscape models can also be extended to include epigenetic effects. Epigenetic mechanisms allow for plastic responses to environmental changes, for example by gene regulation mechanisms [139], and they are frequently heritable through non-genetic inheritance. Some theoretical work has been done to establish that epigenetics can substantially affect evolution on adaptive landscapes [140- 142]. However, epigenetic-genetic interactions are very hard to measure, due to the real-time nature of epigenetic effects and their dependence on environmental conditions. More data will be required for realistic evolutionary simulations encompassing epigenetic-genetic interactions.

## Conclusions

We believe the following three issues will be important for the study of adaptive landscapes in the near future:

(1) New technologies and methodologies must be designed and deployed to efficiently measure diverse, complex, and sizable empirical adaptive landscapes with high resolution. The interdisciplinary nature of this effort will require collaborations between teams of molecular biologists, geneticists, engineers, physicists, theoreticians, and computer scientists.

(2) New methods and analyses must be developed to extract quantitative properties of the topology of landscapes, as well as the evolutionary dynamics that they lead to. Like other fields in which new theoretical and computational methods are developed, this will likely include integration of knowledge from biochemistry, evolutionary theory, epidemiology, and ecology, with methods from quantitative fields such as information theory, probability theory, topology, and statistical physics.

(3) Dissemination of new ideas and insights into diverse fields such as oncology and agriculture will require interdisciplinary teams combining applied biologists and theoreticians.

Adaptive landscapes are a very important factor in evolutionary biology, and as more and more empirical landscapes are measured and published, they will be used to inform our understanding of many biological phenomena, from drug discovery and cancer treatment to community ecology and conservation biology.

## Acknowledgements

The research has been supported in part by ISF 1568/13, an EMBO postdoctoral fellowship (UO) and the Stanford Center for Computational, Evolutionary and Human Genomics (YR). We would like to thank Ohad Lewin-Epstein for his helpful suggestions and review of the manuscript.

## Glossary

Genome: The heritable genetic information of an individual organism.
Gene: A sequence of nucleotides in the genome, the basic functional unit determining hereditary traits.
Locus: The location of a gene on the genome.
Allele: Variant of a specific gene; multiple alleles of the same gene can differ in sequence and function.
Genotype: A specific allele combination.
Phenotype: The physical manifestation of a genotype, including the sum of the organisms’ traits such as metabolism, behavior, morphology, etc.
Evolution: The change in frequencies of different alleles and allele combinations in populations over time.
Natural selection: The sum of the processes that drive changes in frequencies of heritable traits due to the differences in reproductive success these traits induce; for example, differences in growth rates, survival, or fecundity.
Random genetic drift: The sum of the processes that drive changes in frequencies of heritable traits due to random effects, disregarding natural selection; for example, sampling error in small populations.
Recombination: The process in which alleles from two genotypes combine to produce new allele combinations.
Fitness: The contribution of specific genotypes to the next generations, usually given by the reproductive success of individuals carrying these genotypes, which integrates factors such as survival, fecundity, etc.
Mutation: The process in which alleles randomly change to other alleles.
Epistasis: Interactions between different loci such that the phenotype or fitness effect of an allele changing depends on the identities of alleles in other loci.
Sign epistasis: Interactions between loci that change an allele’s contribution to fitness from deleterious to beneficial and vice versa.
Fixation probability: The probability that an allele will take over a population. Usually refers to a rare allele, integrating the deterministic effect of natural selection with the stochastic effect of random genetic drift.

